# Diversity of active viral infections within the *Sphagnum* microbiome

**DOI:** 10.1101/320002

**Authors:** Joshua M.A. Stough, Max Kolton, Joel E. Kostka, David J. Weston, Dale A. Pelletier, Steven W. Wilhelm

## Abstract

*Sphagnum-dominated* peatlands play an important role in global carbon storage and represent significant sources of economic and ecological value. While recent efforts to describe microbial diversity and metabolic potential of the *Sphagnum* microbiome have demonstrated the importance of its microbial community, little is known about the viral constituents. We used metatranscriptomics to describe the diversity and activity of viruses infecting microbes within the *Sphagnum* peat bog. The vegetative portions of 6 *Sphagnum* plants were obtained from a peatland in northern Minnesota and total RNA extracted and sequenced. Metatranscriptomes were assembled and contigs screened for the presence of conserved virus marker genes. Using bacteriophage capsid protein, gp23, as a marker for phage diversity, we identified 33 contigs representing undocumented phage s that were active in the community at the time of sampling. Similarly, RNA-dependent RNA polymerase and the Nucleo-Cytoplasmic Large DNA Virus (NCLDV) major capsid protein were used as markers for ssRNA viruses and NCLDV, respectively. In total 114 contigs were identified as originating from undescribed ssRNA viruses, 22 of which represent near-complete genomes. An additional 64 contigs were identified as being from NCLDVs. Finally, 7 contigs were identified as putative virophage or polinto-like viruses. We developed co-occurrence networks with these markers in relation to the expression of potential-host housekeeping gene *rpb1* to predict virus-host relationships, identifying 13 groups. Together, our approach offers new tools for the identification of virus diversity and interactions in understudied clades, and suggest viruses may play a considerable role in the ecology of the *Sphagnum* microbiome.

**Significance:** *Sphagnum-dominated* peatlands play an important role in maintaining atmospheric carbon dioxide levels by modifying conditions in the surrounding soil to favor its own growth over other plant species. This slows rates of decomposition and facilitates the accumulation of fixed carbon in the form of partially decomposed biomass. The unique environment produced by *Sphagnum* enriches for the growth of a diverse microbial consortia that benefit from and support the moss’s growth, while also maintaining the hostile soil conditions. While a growing body of research has begun to characterize the microbial groups that colonize *Sphagnum*, little is currently known about the ecological factors that constrain community structure and define ecosystem function. Top-down population control by viruses is almost completely undescribed. This study provides insight into the significant viral influence on the *Sphagnum* microbiome, and identifying new potential model systems to study virus-host interactions in the peatland ecosystem.

## Introduction

Peatlands represent one of the most significant biological carbon sinks on the planet, storing an estimated 25% of terrestrial carbon in the form of partially decomposed organic matter (1–3). This accumulation of carbon is achieved through much slower rates of respiration and decomposition than observed in soil, due in large part to the low pH, nutrient-poor, and anaerobic environments created by the dominant moss population (4, 5), of which the genus *Sphagnum* is most prevalent (6, 7). As these environmental conditions appear to favor the growth of *Sphagnum* over vascular plants, primary production is dominated by the moss, which further retards decomposition due to production of antimicrobial compounds such as sphagnic acid (8–10) and sphagnan (11, 12). Despite this, *Sphagnum* and other peat mosses cultivate a diverse, symbiotic microbiome that appears to abate nutritional gaps for the moss and also contribute to the unique biogeochemical characteristics of the peatland ecosystem (13–15). In addition to their value as reservoirs of microbial diversity, the partially decomposed organic matter, known as *Sphagnum* peat, serves as an important economic resource for use in horticulture. As many peat bogs have begun to experience stress due to anthropogenic disturbances (16–18) and possibly climate change (19), the *Sphagnum* microbiome is of interest in peatland conservation and the ecosystem’s services to the surrounding environment.

While there is a growing body of research characterizing the microbial groups that colonize *Sphagnum* (15), little is currently known about the ecological factors that define community structure and ecosystem function. Studies suggest that subtle differences in pH and available nutrients, manipulated by different *Sphagnum* species and strains, create distinct microbial consortia (14, 20, 21). Other observations suggest a more homogenous community (22), highlighting a need for further study. Culture-dependent experiments isolating endophytic bacteria indicate *Sphagnum* cultivates symbionts with abilities that include antifungal activity (20, 23) and nitrogen fixation (14), and that these microbiomes may be passed vertically to the moss progeny (21). Yet while examinations of how environmental conditions and host-microbe symbiotic interactions shape the structure and function of microbial communities, the influence of virus populations on the *Sphagnum* microbiome remains unexplored.

Viruses are the most abundant biological entities on Earth, and central to global ecosystems as they can drive the host evolution through predator-prey interactions and horizontal gene transfer (24). Moreover, viruses can lyse single-celled primary producers and heterotrophs, releasing nutrient elements from the biomass of prokaryotes and eukaryotic protists (25, 26). Viruses may also act as a top-down control on the composition and evenness of microbial communities, targeting hosts that reach higher cell densities, a phenomenon referred to as the “*kill-the-winner*” model (27).

As lab studies of viruses require hosts that can be grown in culture, many environmentally relevant viruses are poorly understood and their representation in reference databases is often skewed. Previous efforts to describe environmental viromes have focused primarily on the sequencing of shotgun or PCR-targeted metagenomes. While these methods have proven powerful, rapidly expanding available reference material for bacteriophage (28, 29), it leaves the considerable diversity of RNA viruses largely untapped (30). Moreover, the common approach of selecting for viruses based on size-exclusion with filters removes many of the Nucleo-Cytoplasmic Large DNA Viruses (NCLDVs, or commonly “giant viruses”) that are also environmentally relevant and phylogenetically informative (31, 32). Metagenomic sequencing also limits observations to virus particles: from these data inferences on viral activity require tenuous assumptions. The advent of high-throughput RNA sequencing offers viral ecologists the opportunity to study active infections in the environment, as DNA viruses only produce transcripts inside a host. Moreover, this approach also captures fragments of RNA virus genomes. When sequencing is of sufficient depth and multiple samples are collected with spatial and temporal variability, these data present an opportunity to develop hypothetical relationships between virus and host markers (33) for subsequent in lab testing.

In this study, we analyzed metatranscriptomes from the microbial community inhabiting the vegetative portion of *Sphagnum fallax* and *S. magellanicum* plants in Northern Minnesota, with the goal of describing active viral infections within the *Sphagnum* microbiome. Using marker genes conserved within several viral taxa, we identified an active and diverse bacteriophage population, largely undescribed in previous studies. We also identified ongoing infections by a diverse consortium of “giant” viruses and potentially corresponding virophage/polinton-like viruses (hereafter referred to as virophage), including several giant viruses closely related to the recently discovered Klosneuviruses (34). Finally, a number of novel positive-sense single-stranded RNA viruses, some of which assembled into near complete genomes, were observed. With this information in hand we developed statistical network analyses, correlating co-expression of viral marker genes with housekeeping transcripts from potential hosts. The resulting observations propose several virus-host pairings that, moving forward, can be tested in a laboratory setting. Together, these results demonstrate new potential model systems to study virus-host interactions in the peat bog ecosystem, and provide insight into the significant viral influence on the *Sphagnum* microbiome.

## Results

### Identification of resident phage populations

To identify active virus populations in the *Sphagnum* phyllosphere, we obtained *S. fallax and S. magellanicum* plant matter samples (3 from each species) from peatland terrariums as a part of the Spruce and Peatlands Under Changing Environments (SPRUCE) project for metatranscriptomic sequencing. Across all six *Sphagnum* phyllosphere samples, 33 contigs were identified as transcripts encoding major capsid protein *(gp23)* originating from bacteriophage, while only 6 contigs were identified using three other marker genes. Concurrent with this, more reads mapped to *gp23* contigs than to the other marker genes combined, the most abundant of which were three ribonucleotide reductase contigs.

Of the 33 contigs, 18 were assigned to the *Eucampyvirinae* subfamily with *Campylobacter* viruses CP220 and PC18, while the rest were spread amongst the other Myovirus taxa, predominantly the *Tevenvirinae* (Fig 1). SS4 contig 77559 was the most abundant, with consistently high expression across all samples, whereas other contigs dominated just one or two samples. Of the 6 contigs identified using the other 3 viral marker genes, one was identified as a potential *gp20* homologue, originating within *Myoviridae* with *Clostridium* virus phiCD119 as the closest relative (SFig 1). Two contigs were identified as *recA* contigs, likely originating in myovirus and siphovirus relatives (SFig 2), while the remaining three contigs were identified as ribonucleotide reductase transcripts (SFig 3).

**Figure 1:**
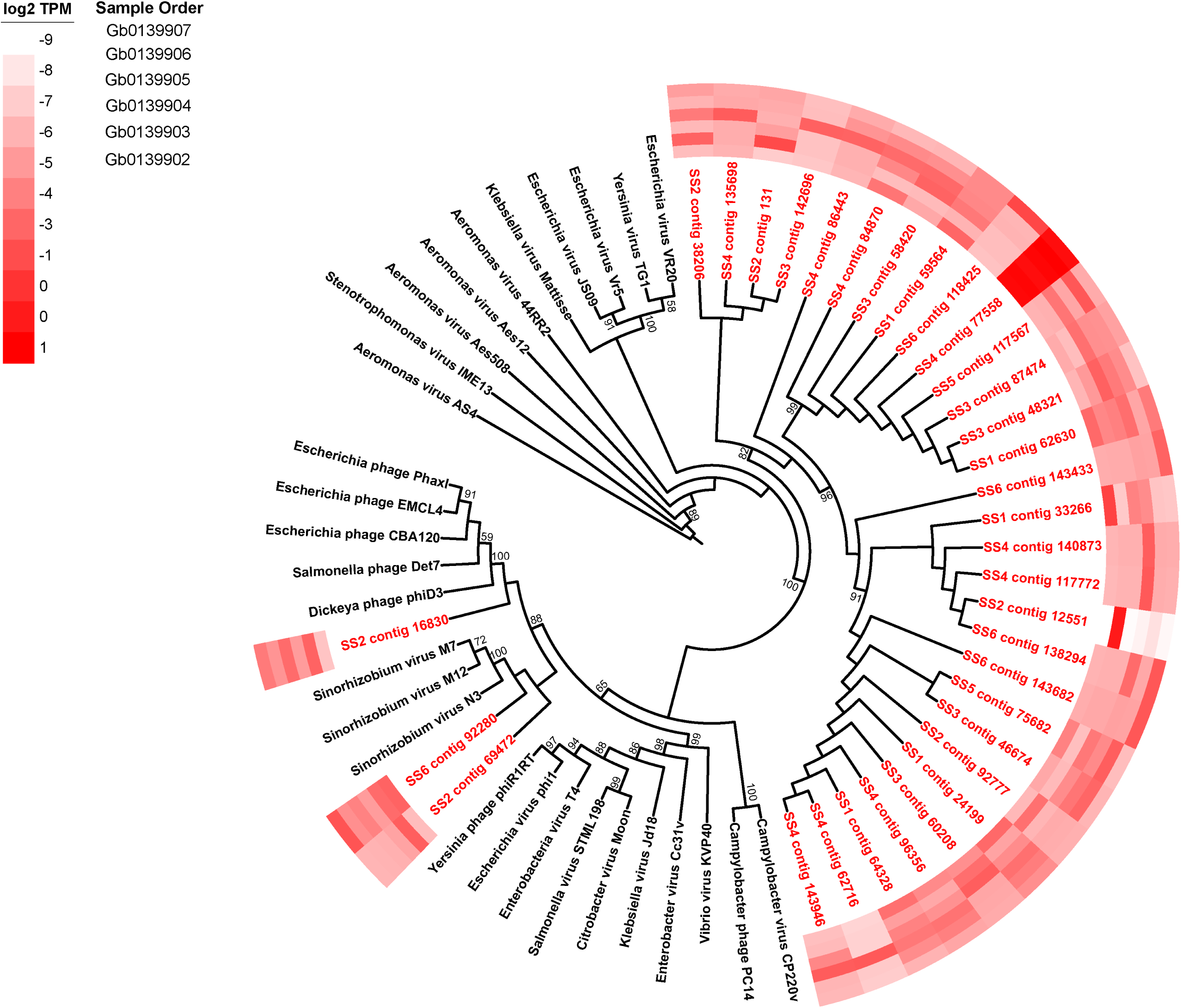
Phylogenetic placement of identified phage major capsid protein contigs (red) on a Myovirus *gp23* maximum likelihood reference tree (references in black). Node support (aLRT-SH statistic) > 50% are shown. Contigs are shown with their abundance (log2 transformed TPM) in a heatmap surrounding the tree. Sample order on the heatmap is provided in the inset.

### Single-stranded RNA virus diversity and abundance

Within our samples, 114 contigs originated from RNA viruses, the majority of which belonged to the currently unassigned *Barnaviridae* and Astrovirus-like families (Fig 2). Additionally, a large number of *Picornaviruses* were observed, most of which were closely related to the unclassified marine *Aurantiochytrium* single-stranded RNA virus, and *Secoviridae* plant viruses. Lastly, several contigs were closely related to the *Nidovirales* clade, which generally infect animal species.

**Figure 2:**
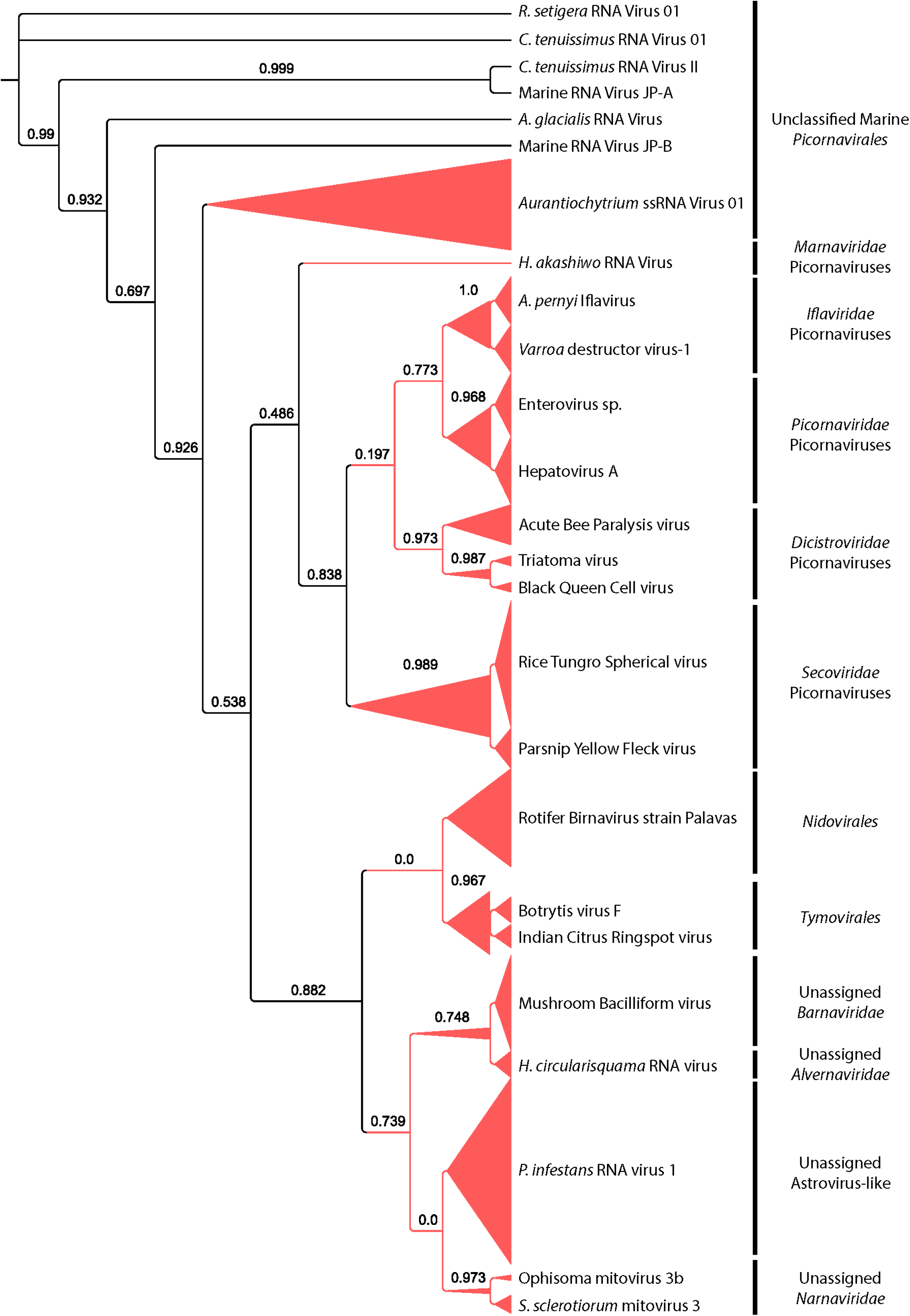
Phylogenetic placement of identified ssRNA virus RNA-dependent RNA polymerase contigs on maximum likelihood reference tree. Branch width represents the number of contigs placed on the reference branch. Node support (aLRT-SH statistic) >50% are shown.

Among these, 22 contigs were found to be near complete ssRNA virus genomes (based on gene content and size), encoding multiple viral genes in addition to RDRP. Gene regions were identified and annotated using the NCBI conserved domain and PFam HMM search tools, and the full-length RDRP sequence was used to construct a maximum likelihood phylogenetic tree (Fig 3). Of the partial ssRNA genomes that were assembled, 2 were missing the conserved Rhv structural genes, while one was missing a RNA virus Helicase. The majority of these contigs fall under the *Picornavirales* order, which also included the most complete viral genomes. As was observed with the shorter RDRP contigs above, most of the *Picornavirales* contigs were most closely related to the unclassified marine species, or members of the family *Secoviridae* clade, whose membership includes the Parsnip yellow fleck virus. A number of partial *Picornavirus* genomes were also identified as members of the family *Dicistroviridae*. Outside the *Picornavirales*, most contigs clustered closely with the unassigned Astrovirus-like *Phytophthora infestans* RNA virus. To determine the relative abundance of different RNA virus genomes in the peat bog samples, we mapped reads back to contigs and calculated transcripts per million (TPM) values to account for contig length and library size. The most abundant contig across all samples was SS4 contig 3964, which was most closely related to the Rotifer birnavirus. All other contigs appear to be abundant prominently in one or two samples, and absent or in low abundance in the others, with no patterns of abundance apparent.

**Figure 3:**
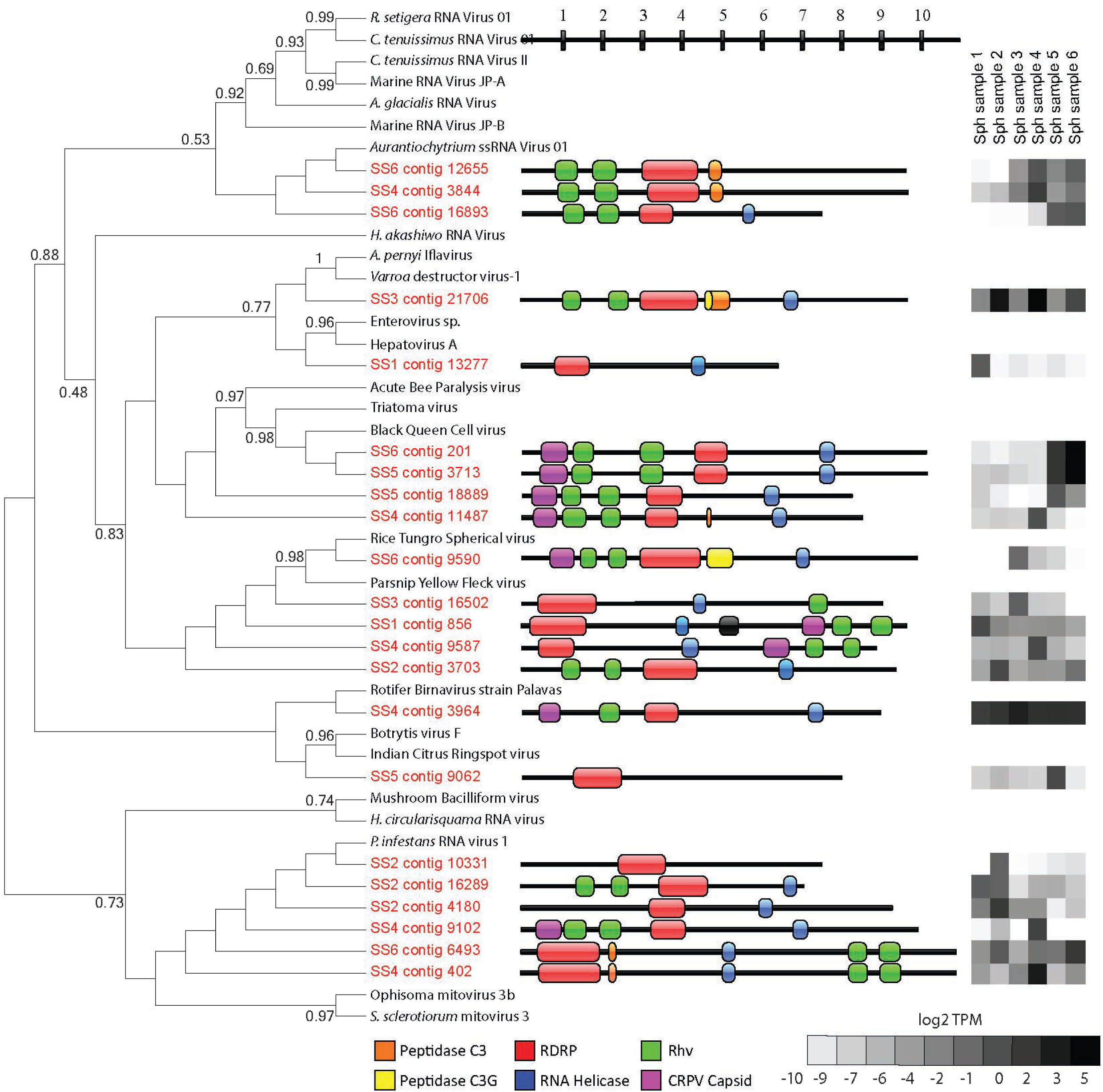
Phylogeny, genome architecture, and abundance of partial ssRNA virus genomes. Tree represents phylogenetic placement of RDRP gene regions from partial ssRNA virus genome contigs (red) on a maximum likelihood reference tree (references in black). Node support (aLRT-SH statistic) >50% are shown. Center panel represents genome architecture determined by conserved domain search and ORF prediction. Length of contigs and gene regions is measured in kb. Heat map in right panel shows abundance of reads mapped to partial genome contigs in log2 TPM from each of the 6 metatranscriptome libraries.

### Giant viruses and virophage in Sphagnum microbiome

Of the 10 gene markers tested to identify Nucleo-Cytoplasmic Large DNA Viruses (NCLDVs), only the giant virus major capsid protein (MCP) was detected in the metatranscriptome. 64 contigs were observed with homology to MCP, representing every known group of NCLDVs (Fig 4). Out of the 64 MCP contigs, 46 were placed within the *Mimiviridae* taxa. Most contigs (25) closely aligned with the recently discovered Klosneuviruses, with the Indivirus and Catovirus representing the most diversity in these samples. The next most abundant group were the “extended *Mimiviridae”* (7 contigs), species with known similarity to Mimiviruses but that infect eukaryotic algae. Six contigs phylogenetically were similar to the *Asfarviridae*, here represented by the African swine fever Virus. Potential relatives of the giant virus outliers, Pandoravirus and Pithovirus, were not observed (due to methodological limitations), and the *Iridoviriae* were poorly represented (1 contig). Using the virophage MCP and packaging ATPase as markers, we identified 7 contigs as transcripts originating in putative virophage or polinton-like viruses, all of which were phylogenetically placed amongst isolates identified from freshwater ecosystems (Fig 5).

**Figure 4.**
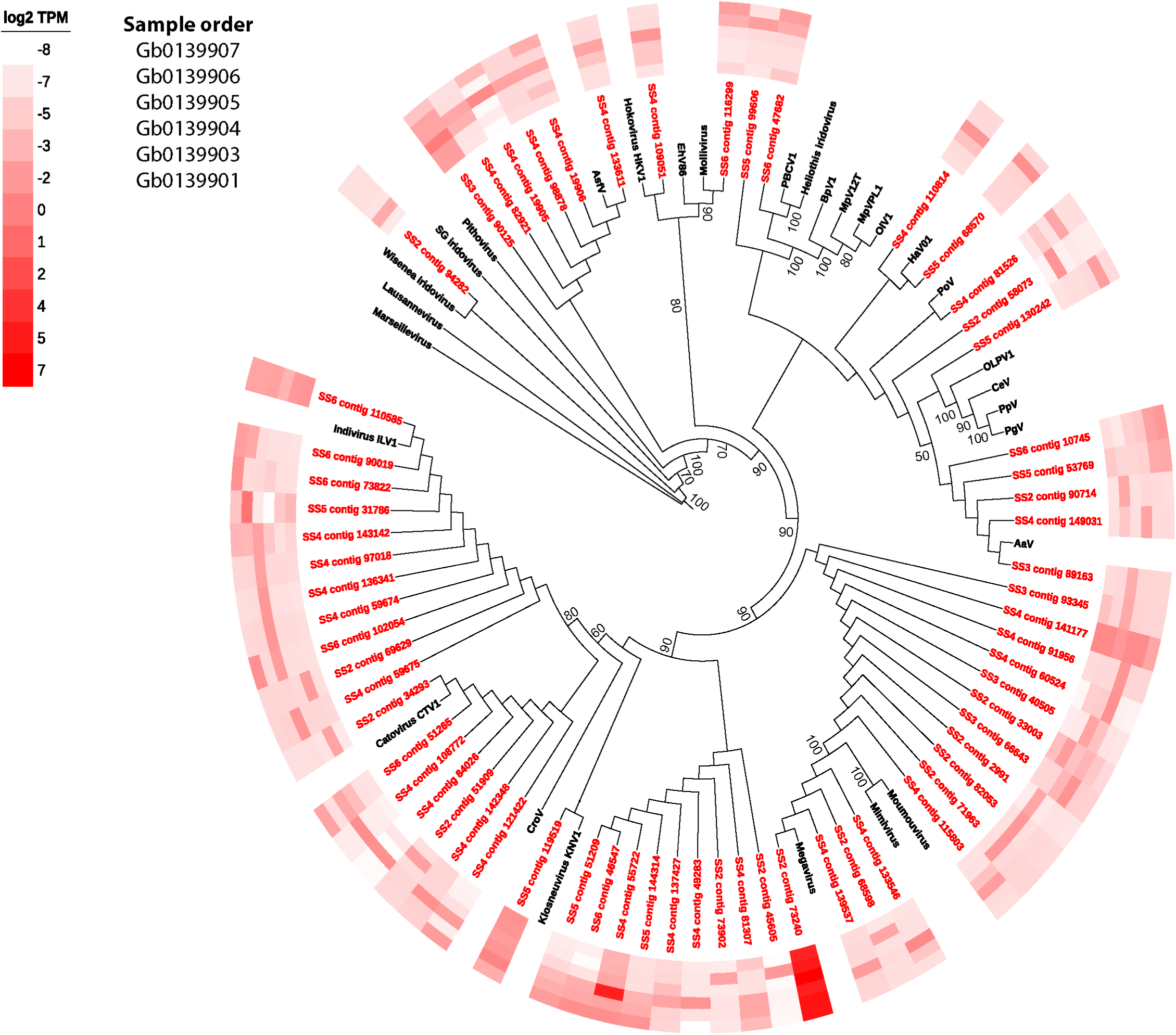
Phylogenetic placement of identified NCLDV major capsid protein contigs (red) on a maximum likelihood reference tree (references in black). Node support (aLRT-SH statistic) >50% are shown. Contigs are shown with their abundance (log_2_ transformed TPM) in a heatmap surrounding the tree.

**Figure 5:**
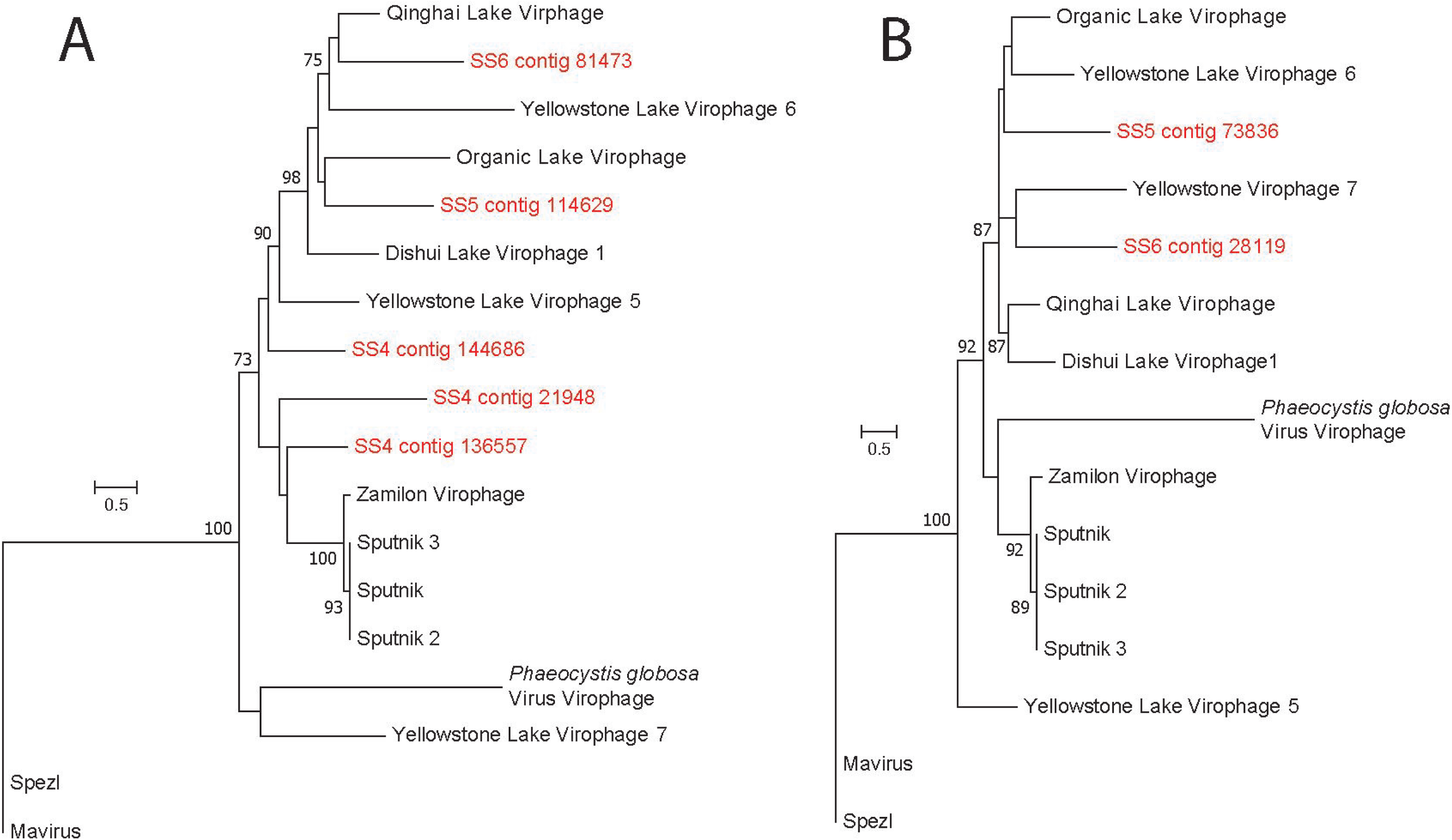
Phylogenetic placement of identified virophage A.) major capsid protein and B.) ATPase contigs (red) on a maximum likelihood reference tree (references in black). Node support (aLRT-SH statistic) >50% are shown.

As was observed with the other major viral taxa described, the majority of contigs were most abundantly expressed in one or two samples and present at very low levels in the rest. The most abundant NCLDV-MCP contig in the samples was SS2 contig 73240, most closely related to *Megavirus chilensis*, which was the most highly expressed giant virus contig across all samples. Four other contigs (SS6 contig 110585, SS4 contigs 55722 and 141177, and SS5 contig 119519) were highly expressed across all six samples.

### Prediction of virus-host pairs

By comparing and correlating expression of virus marker genes to *rpb1* expression from cellular organisms, we endeavored to predict potential virus-host groups in the *Sphagnum* phyllosphere. Fig 6 shows statistically robust networks containing at least one virus and one host, where co-occurrence and correlation were observed in more than one sample. A total of 13 virus-host groups were detected, spread across the major viral taxa detected in this dataset. We note that no networks containing the virophage/polinton-like viruses emerged. Four relationships were predicted from bacteriophage *gp23* abundance, the simplest of which was a *Tevenvirinae phage-Metazoa-Rhizaria* group with moderate correlations (Fig 6a). The other 3 relationships are more complicated, containing multiple potential hosts and, for the largest predicted group, multiple virus transcripts. The majority of potential hosts in these groups were identified as eukaryotic, with only one putative bacterium and two archaea. Correlation coefficients for the phage-prokaryote clusters were lower than was observed in the other major viral taxa, with low to moderate correlations between viruses and bacteria.

**Figure 6:**
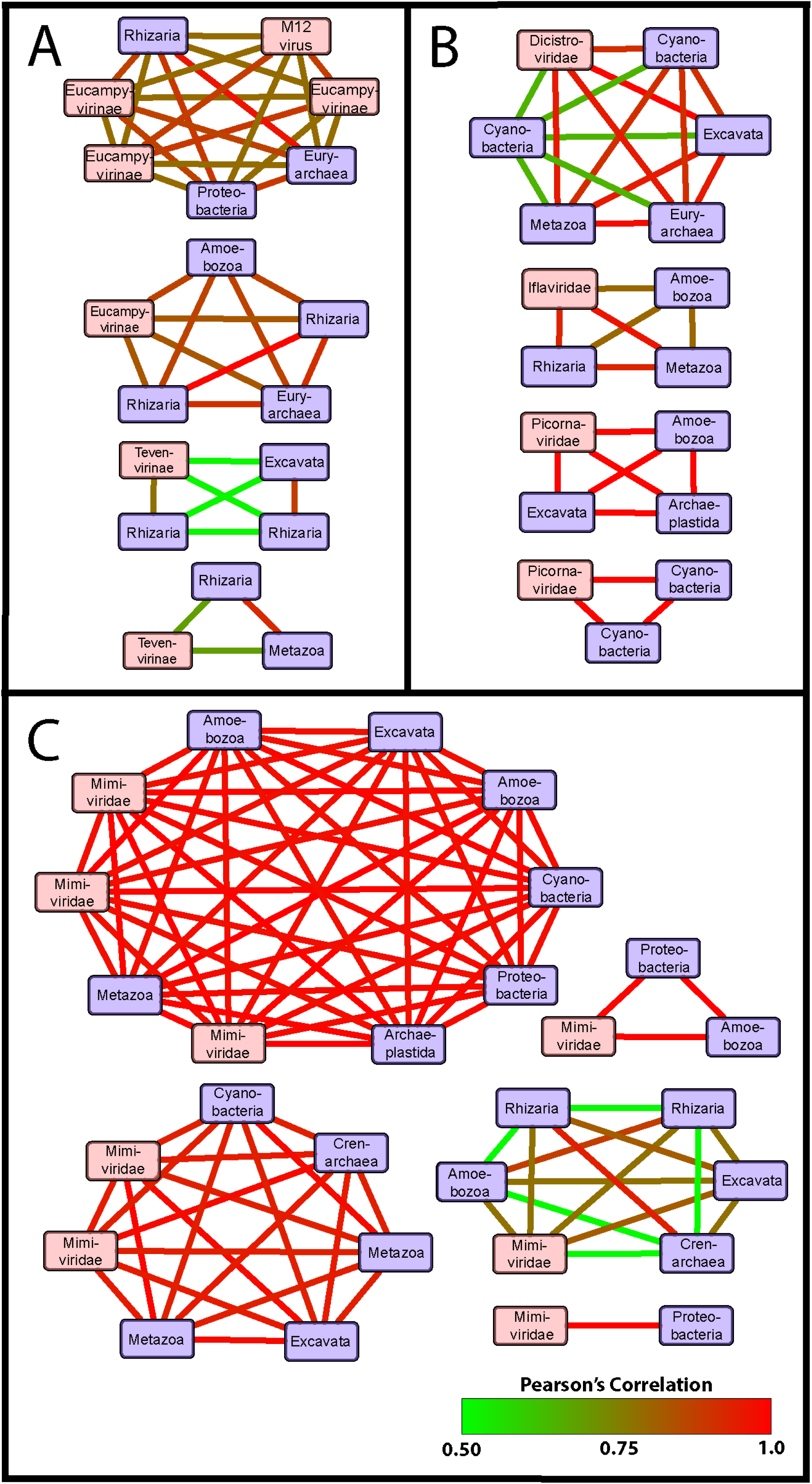
Correlation co-occurrence network analysis of conserved viral gene and host RNA polymerase expression for A.) bacteriophage (Gp23), B.) ssRNA viruses (RDRP), and C.) NCLDVs (NCLDV MCP). Nodes in red represent virus contigs and blue nodes represent potential hosts. Nodes are connected by edges colored according to the Pearson correlation coefficient values between to contigs. Only relationships with contigs expressed in more than one sample are shown.

We observed 4 predicted RNA virus-host clusters, all of which contained multiple hosts grouped with a single virus (Fig 6b). Most of the predicted hosts appear closely related to eukaryotic single-celled protists, within the *Excavata* and *Rhizaria* supergroups. Correlation coefficients observed in these relationships are generally higher than observed in the phage-host clusters. The 5 predicted NCLDV-host clusters (Fig 6c) were the most highly correlated and complex. Predicted hosts were highly varied, ranging from diatoms to animals, though all virus members were placed either within *Mimiviridae* or the extended Mimivirus group. MCP contigs originating in close relatives of the recently discovered Klosneuviruses are present in both the 7- and 10-member clusters, in addition to a pair of contigs most closely related to *Aureococcus anophagefferens* Virus (AaV). An additional 15 statistically significant clusters across all three viral taxa were observed where the virus and host were present in only one sample (not shown).

## Discussion

Understanding the virus burden on microbial communities in ecologically-rich ecosystems is an important step forward in resolving their function and predicting how they might respond to various drivers of ecosystem scale change. In the present study we used metatranscriptomes to describe the diversity and activity of the resident virus populations in a peat moss (*Sphagnum*) microbiome. We identified previously undescribed virus activity from multiple taxa, most of which are poorly represented in either the literature or reference sequence databases. We used read mapping to quantify the relative abundance of active viral infections. Lastly, we compared expression of viral transcripts to that of potential hosts, using a correlation co-occurrence networks approach (33) to predict putative hosts for the observed virus populations. Together, our results suggest that the *Sphagnum* phyllosphere represents a significant and largely untapped source of virus diversity and activity. Viruses were highly active across all samples, with some individual viruses exhibiting abundant activity in single samples, while others were more pervasive. Given that our observations were based on RNA sequencing data, they do not represent a full accounting of the virus particles present in the community. However, metatranscriptomic data, allows us to distinguish virus populations active at the time of sampling. In addition, as viruses only transcribe their genes during infection, virus and host transcripts are expected to co-occur, and it is possible that the abundance of transcripts (at least for DNA viruses) could be used to predict natural hosts of viruses observed in the ecosystem which can be tested in a laboratory or field setting. Ultimately, this study identifies from within a complex community a number of candidate virus-host model systems for future study.

### Viral diversity and activity in Sphagnum plants

As viruses lack a universal genetic marker like the bacterial 16S rRNA gene, we opted to screen metatranscriptome assemblies for genes previously demonstrated to be largely or wholly conserved amongst individual viral taxa. Within the expanded and diverse genetic potential of giant viruses, only a handful of genes are currently conserved amongst all members (32, 35) and these, in addition to several markers conserved amongst a large portion of giant viruses were used to identify activity in the *Sphagnum* phyllosphere. Out of the 10 genes used to screen the metatranscriptomes, we only MCP transcripts. This is not surprising given the number of capsid proteins needed for viral assembly: indeed this transcriptional pattern was previously observed in both cultures (36) and marine systems by Moniruzzaman *et al*. (2017). It should be noted that the RNA-seq dataset used in those studies was poly-A selected, enriching for eukaryotic transcripts, and thus coverage of eukaryotic virus gene expression would be much higher than in the *Sphagnum* metatranscriptome. That we observed MCP expression in abundance suggests a significant number of infections occurred at the time of sampling. While the magnitude of giant virus diversity in *Sphagnum* dominated ecosystems is, to our knowledge, completely unexplored, the richness observed here is considerably larger than expected compared to better documented systems. 64 distinct MCP genotypes were identified in the *Sphagnum* phyllosphere metatranscriptomes, which is high when compared to one recent survey that identified 30 novel MCP transcripts from multiple environmental datasets (37), and another which observed 107 NCLDV sequences in 16 publicly available environmental metagenomes of comparable sequencing depth isolated from different ecosystems (38). Most of the MCP contigs identified were placed in clusters around a small number of virus relatives, highlighting the under-sampled diversity of giant viruses in the literature, poor representation in reference databases, and the considerable diversity present in *Sphagnum* peat bogs. The significant giant virus diversity observed here implies a corresponding eukaryotic richness that is also under-described (39). Additionally, a series of virophage transcripts were detected, indicating a significant response to infections by giant viruses in the system. Many of these are phylogenetically grouped with the polintoviruses, transposable elements that produce virion particles that can exploit the replication machinery of actively infecting giant viruses to reproduce, often at the expense of the giant (40, 41). These observations suggest that while an active picoeukaryotic population may persist, mortality mechanisms beyond grazer-driven losses are at play and likely important to carbon flow in the system.

The use of RNA-seq presents a unique opportunity to capture the genomic material of RNA viruses that is lost in metagenomic sequencing. As such, RNA virus representation in sequencing databases and the literature is largely constrained to culture-based studies. All known RNA viruses require a functional RNA-dependent RNA polymerase (RdRP) to copy their genome inside the host cell, a function exclusive to viruses, making it a highly specific marker for RNA virus discovery (42, 43). Recent attempts to use metatranscriptomes to describe environmental RNA viruses have proven successful, not only identifying marker gene fragments in datasets, but assembling complete and near-complete genomes (33, 43). The diversity and composition of RNA virus populations in *Sphagnum* peatlands is largely unknown: it is currently limited to the small group of RNA-DNA hybrid chimeric Cruciviruses (44). Here, as was observed with the giant viruses, most RNA virus contigs were placed in clusters with a single represented species, suggesting a significant degree of uncharacterized diversity. This is not entirely surprising, as RNA viruses are expected to make up as much as half of the virus particles in the Earth’s oceans, and yet they are almost as poorly understood and represented in sequencing databases as giant viruses (30). Similarly, we assembled and identified 22 near-complete RNA virus genomes, where completeness was determined primarily by size and the presence of the 6 core genes. As there are currently only 265 sequenced genomes within the *Picornavirales*, most of which grouped within the *Picornaviridae*, this represents a sizeable addition to the known diversity of ssRNA viruses. This is especially true for the unassigned and unclassified taxa, and establishes a strong foundation for future efforts to describe RNA virus populations in *Sphagnum*.

Description of bacteriophage populations in *Sphagnum* peatlands is currently limited to the ssDNA viruses of the *Microviridae* (45) and *Caudovirales* (46) observed in metagenomics data, though it appears that phage are the most abundant biological entities in the *Sphagnum* phyllosphere (46). Given this, and the dominance of bacteria in the *Sphagnum* microbiome as previously described (15), the relatively low abundance of active bacteriophage in our samples was a surprise. Marker genes to identify bacteriophage were chosen based on their conservation across phage taxa and their success in other environmental datasets. Gp20 (phage portal protein) and Gp23 (major capsid protein) have been shown previously to be highly conserved and effective for phylogenetic assignment of members of the *Myoviridae* (47–49). RecA is conserved across all three bacteriophage taxa and could illuminate lysogeny, and ribonucleotide reductase (RNR) has been used as an effective marker for screening novel viruses from marine sequencing datasets (50). As such, we identified 39 bacteriophage contigs using these markers, 33 of which were from Gp23. This may represent a similar phenomenon as MCP in the giant viruses above, where transcripts encoding structural proteins are much more abundant than other genes and sequencing lacked the depth to detect them. For the purpose of discovering novel phage species, DNA sequencing through metagenomics may prove more successful.

### Virus-host predictions

Future study of viral dynamics in peatlands will require the establishment of model virus/host pairs for *in vitro* experimentation and *in situ* tracking. While culture-based techniques can yield model systems, it is not always clear whether the isolated organisms are environmentally relevant. In order to address this, we attempted to use statistical methods to propose virus/host pairs as potential future model systems based on their cooccurrence in samples and the correlation of their abundance. As viruses produce transcripts only when actively infecting a host, positive correlation and co-occurrence between virus and host transcripts is expected, and might be used to predict host-virus relationships, provided an appropriate transcriptional proxy for growth and activity is available (33). In this study, we used the eukaryotic RNA-polymerase gene *rpb1* as a marker for abundance and activity in potential hosts, as it is conserved amongst all eukaryotic organisms, is phylogenetically informative, and has been previously described as one of the more consistently expressed eukaryotic genes in marine systems, scaling well with the activity of the organism (51), though the stability of its expression has not been evaluated in terrestrial ecosystems. We used NCLDV MCP abundance as a proxy for giant virus production, Gp23 for phage production, as transcription is necessary for the assembly of new virus particles and transcript abundance in some appears to be closely linked to viral replication. We also used RdRP as a proxy for RNA virus production, acknowledging the caveat that we cannot distinguish between abundance of free virus particles and active infections (33).

Correlation and co-occurrence matrices, clustered into groups by similarity and tested with the SIMPROF permutation test, yielded 13 predicted groups of viruses and hosts. For ssRNA and giant viruses, several of the networks produced in the analysis included multiple bacterial and archaeal sequences picked up in the RNA polymerase screen. As we have no reason to believe bacterial species are infected by NCLDVs or Picornaviruses, it is likely these predictions represent a confounding relationship between prokaryotes and potential eukaryotic hosts, observed in network analyses for all three viral taxa described here, where a beneficial interaction results in an indirect correlation with viral infection. Indeed, previous use of this method in marine systems showed a similar phenomenon, where an algal Mimivirus and a known host were grouped with a fungal species and another virus (33). Even after the consideration of bacterial species within the predicted groups, some remain complicated with multiple viruses and potential eukaryotic hosts, which may be explained by a broader host range amongst giant viruses enabled by the expansion of genetic material and increased independence from host machinery. Similar relationships were observed amongst RNA viruses, though these are more tenuous, as we are unable to distinguish whether sequencing reads originated transcripts or genomic material.

All together, we have identified a considerable amount of viral diversity from several major viral taxa active within a poorly understood microbial ecosystem. As they were identified from transcript sequencing data, the viruses described here likely only represent a fraction of the whole virus community, which may be elucidated through further culture-independent work. We have also used transcript abundance within a statistical framework to predict several host-virus relationships which can be sought out and tested in culture. These results establish an important and much needed foundation for future research into the microbial ecology in *Sphagnum* peat bogs.

## Materials and Methods

### Sample collection and Survey of Environmental Conditions

Triplicate individual plants of *Sphagnum magellanicum* and *Sphagnum fallax* were collected on August 2015 from the SPRUCE experiment site at the S1 bog on the Marcell Experimental Forest (U.S. Forest Service, http://mnspruce.ornl.gov/). The S1 Bog is an acidic and nutrient-deficient ombrotrophic *Sphagnum-dominated* peatland bog (surface pH≤4.0) located approximately 40 km north of Grand Rapids, Minnesota, USA (47°30.476’ N; 93°27.162’ W; 418 m above mean sea level) (52–54). To characterize the *Sphagnum* virome, *Sphagnum* samples were collected as previously described (54). Only green living plants were sampled: samples focused on the capitulum plus about 2-3 cm of green living stem. B *Sphagnum* stems (phyllosphere) were cleaned from unrelated plant debris, and frozen immediately on dry ice. Frozen samples were overnight shipped to the Georgia Institute of Technology for RNA extraction.

### RNA Extraction and Sequencing

One gram of *Sphagnum* phyllosphere tissue was ground with a mortar and pestle under liquid nitrogen. The fine powder was transferred to 10 extraction tubes and total RNA isolated using the PowerPlant RNA Isolation Kit with DNase according to the manufacturer’s protocol (MoBio Laboratories, Carlsbad, CA, USA). DNA-depleted RNA was quantified using the Qubit RNA HS Assay Kit (Invitrogen, Carlsbad, CA, USA) and quality was assessed on the Agilent 2100 BioAnalyzer using the Agilent RNA 6000 Pico Kit (Agilent Technologies). Additionally, the absence of DNA contamination was confirmed by running a polymerase chain reaction using universal bacterial 16S rRNA primers 515F and 806R. Finally, RNA samples without detectible DNA contamination and exhibiting an RNA integrity number (RIN) > 6 were pooled. Extracted total environmental RNA samples were was sent on dry ice to the Joint Genome Institute (JGI) facilities for meta-transcriptomes libraries construction and sequencing. All protocols employed were standard JGI protocols Ribosomal RNA subtraction from total environmental RNA was completed using the Ribo-Zero rRNA Removal Kit (Illumina, San Diego, CA). rRNA depleted environmental RNA were used to construct paired end metatranscriptomes libraries using TruSeq kit and sequenced on the Illumina HiSeq2000 platform at the JGI facilities using a single-end 250bp flow cell.

### RNA-seq Data Processing

Raw sequences (see Supplementary Table 2) were downloaded from the Department of Energy Joint Genome Institute server and processed using the CLC Genomics Workbench v. 10.0.1 (QIAGEN, Hilden, Germany). Reads below a 0.03 quality score cutoff were removed from subsequent analyses, and the remaining reads were trimmed of any ambiguous and low quality 5’ bases. Samples were subjected to a subsequent *in silico* rRNA reduction using the SortmeRNA 2.0 software package (55). Filtered reads were *de novo* assembled with cutoffs of 300 base minimum contig length and average coverage of 2, leaving a total of 705,526 contigs across all samples.

### Screening Assemblies for Marker Genes

Marker genes to identify bacteriophage were chosen based on their conservation across phage taxa and their success in other environmental datasets. Gp20 (phage portal protein) and Gp23 (major capsid protein) have been shown previously to be highly conserved and effective for phylogenetic assignment of members of the *Myoviridae* (47–49). RecA is conserved across all three bacteriophage taxa and could illuminate lysogeny, and ribonucleotide reductase (RNR) has been used as an effective marker for screening novel viruses from marine sequencing datasets (50). To identify contigs specific to the <u>N</u>ucleo<u>C</u>ytoplasmic <u>L</u>arge <u>D</u>NA <u>V</u>irus (NCLDV) clade, contig libraries were screened for the presence of 10 genes previously identified as core NCLDV genes as previously described (33). Briefly, contig libraries were queried against Nucleo-Cytoplasmic Virus Orthologous Groups (NCVOG) protein databases for each of the following 10 marker genes in a Blastx search with a minimum e-value cutoff of 10^−3^: A32 virion packaging ATPase (NCVOG0249), VLFT-like transcription factor (NCVOG0262), Superfamily II Helicase II (NCVOG0024), mRNA capping enzyme (NCVOG1117), D5 helicase-primase (NCVOG0023), ribonucleotide reductase small subunit (NCVOG0276), RNA polymerase large subunit (NCVOG0271), RNA polymerase small subunit (NCVOG0274), B-family DNA polymerase (NCVOG0038), and major capsid protein (NCVOG0022). Resulting hits were then queried against the NCBI refseq protein database (56) and only contigs with top hits to virus genes were maintained for subsequent analyses. A similar method was used to identify virophage transcripts, where the virophage major capsid protein and packaging ATPase genes were used as markers.

Contigs derived from ssRNA viruses were identified by screening the contig library for RNA-dependent RNA Polymerase (RDRP), a distinctive and wholly conserved RNA virus gene and a strong phylogenetic marker (57). A BLAST database of RDRP sequences was downloaded from the pfam database (58) under code pf00680. Contigs were aligned using Blastx with a minimum evalue of 10^−4^. Hits were queried against the NCBI refseq protein database and only hits to viral RDRP genes were retained for downstream analyses.

To identify RNA virus genome fragments, contig libraries were screened as described above using the following core set of genes observed in RNA viruses: CRPV capsid (Pfam 08762), VP4 (Pfam 11492), RdRP (Pfam 00680), Peptidase C3 (Pfam 00548), Peptidase C3G (Pfam 12381), Rhv (Pfam 00073), and RNA Helicase (Pfam 00910). BLAST databases for core RNA virus genes were constructed from reference sequences downloaded from pfam. Query sequences were then cross-referenced to identify contigs with hits to multiple RNA virus core genes. Only contigs > 1000 bases with at least one viral RDRP region were retained for further analysis. ORFs were predicted on these putative partial genomes using the CLC Genomics Workbench. Features on the partial genomes were predicted using the Pfam HMM domain and the NCBI Conserved Domain Database searches (59, 60). Genome architecture was visualized using the Illustrator for Biological Sequences (IBS) software package (61).

### Phylogenetic Analysis

Reference sequences for viral marker genes and Rpb1 were downloaded from the InterPro and RefSeq databases (STable 1) (62). Reference sequences were aligned using the MUSCLE alignment algorithm (63) in the MEGA v7.0.26 software package (64). Maximum likelihood phylogenetic trees were constructed in PhyML (65) with the LG substitution model and the aLRT SH-like likelihood method. Putative viral and Rpb1 contigs assembled from the metatranscriptomes were translated into proteins according to the reading frame of the top BLAST hit. Translated proteins were placed on the reference trees in a maximum likelihood framework in pplacer (66). Trees with abundance data were visualized using the iToL web interface (67).

### Statistical Analysis

Quality filtered and trimmed reads were stringently mapped to the selected contigs (0.97 identity fraction, 0.7 length fraction) in CLC Genomics Workbench 10.0.1. Expression values were calculated as a modification of the transcript per million (TPM) metric. Read counts were normalized by contig length in kb to determine the reads per kilobase (RPK) values for every contig within each library. These RPK values were then summed and divided by 1 million, to determine the sequencing depth scaling factor for each library. TPM for a contig was calculated by dividing its RPK value by the scaling factor for the library.

Expression values for contigs were imported into the PRIMER7 (68) statistical software package and log2 transformed. Expression values from each contig were correlated (Pearson’s rho) to one another and statistically grouped by co-occurrence using group average hierarchical clustering. The SIMPROF test (69) was used to determine the statistical significance level of resulting clusters (alpha = 0.05, 1000 permutations). Statistically significant clusters with at least one viral contig, one *rpb*1 contig and less than 10 total members were visualized and annotated in Cytoscape 3.5.1 (70).

### Accession Numbers

Full RNA-seq libraries have been made publicly available on the JGI website under accession number Gp0146911.

## Acknowledgements

Research sponsored by the *Laboratory Directed Research and Development* Program of Oak Ridge National Laboratory and the *Joint Directed Research and Development Program* of the University of Tennessee. Support for the SPRUCE experimental site is from the U.S. Department of Energy, Office of Science, Office of Biological and Environmental Research. Oak Ridge National Laboratory is managed by UT-Battelle, LLC, for the U.S. Department of Energy under contract DE-AC05-00OR22725. Support at UT was received from the *Kenneth & Blaire Mossman Endowment* to the University of Tennessee (SWW).

